# Chikungunya virus infection impairs osteogenic differentiation of bone marrow-derived mesenchymal stem cells

**DOI:** 10.1101/780791

**Authors:** Enakshi Roy, Wen Shi, Bin Duan, St Patrick Reid

**Affiliations:** Department of Pathology & Microbiology, University of Nebraska Medical Center, Omaha, NE 68198-5900, USA; Mary & Dick Holland Regenerative Medicine Program, Division of Cardiology, Department of Internal Medicine, University of Nebraska Medical Center, Omaha, NE 68198-5900, USA

**Author notes:** Corresponding authors- St Patrick Reid,; Bin Duan.

**Keywords:** Chikungunya virus, mesenchymal stem cells, osteogenic differentiation

## Abstract

Chikungunya virus (CHIKV) is a positive-sense, single-stranded RNA virus, belonging to the genus alphavirus in the family *Togaviridae*. The virus is spread by the *Aedes* species (sp.) mosquitoes in tropical and subtropical regions of the world. CHIKV causes Chikungunya fever (CHIKF), where the acute stage of infection is characterized by high fever, headache, rash, and polyarthralgia. In 30-40% of cases, patients develop a chronic stage with debilitating joint pain persisting for months to years imposing a burden on the population in terms of disability adjusted life years (DALY). Presently, no vaccines or treatment options are available for this infection. Prior investigations reveal that CHIKV infection is associated with bone pathology; however, the molecular mechanism underlying CHIKV-induced bone pathology remains poorly defined. Studies show that disruption of osteogenic differentiation and function of bone marrow-derived mesenchymal stem cells (BMMSCs) can lead to bone pathologies. However, to date pathogenesis of CHIKV infection in this context has not been studied. In the current study, we investigated the susceptibility of BMMSCs to CHIKV and studied the effect of infection on BMMSCs-derived osteogenic cells. To our knowledge, for the first time we report that CHIKV can productively infect BMMSCs. We observed a decrease in the intracellular and extracellular alkaline phosphatase (ALP) activity and reduction in calcium phosphate deposition in infected cells compared to mock-infected control. Thus, we conclude that CHIKV infects BMMSCs and disrupts function of osteogenic cells.

**Importance:** Although studies have shown association of bone pathology and CHIKV infection, the pathogenesis of infection causing altered bone homeostasis is not fully understood. Here, we demonstrate for the first time that BMMSCs are susceptible to CHIKV infection. Furthermore, we observe that infection causes disruption in the function of BMMSC- derived osteogenic cells. Impaired function of these osteogenic cells will likely lead to a disruption in bone homeostasis and in part, provides a mechanism for the observed bone pathology associated with CHIKV pathogenesis.

## Introduction

Chikungunya virus (CHIKV) is a positive-sense, single-stranded RNA virus belonging to the *Togaviridae* family and alphavirus genus (1) (2). Since the mid-1900s, there have been outbreaks of CHIKV infection in Africa, Asia, and the Indian and Pacific Ocean region, with few reported cases within Europe (3). Beginning in 2013, there have been increasing numbers of infections in the Americas, in part due to travel from affected region (4). CHIKV is transmitted by *Aedes* species (sp). Mosquitoes. CHIKV infection in the tropical and subtropical regions of the world can lead to Chikungunya fever (CHIKF) (5). CHIKF is characterized by a self-limiting acute stage, with symptoms of fever, rash and arthralgia which lasts for 1-2 weeks (6). In 30-40% of cases, infected individuals develop an incapacitating chronic arthritic stage which may persist for months to years, thereby imposing a burden on the population in terms of disability adjusted life years (DALY) (7–10)

Recent studies identified bone lesions in the joints of CHIKV infected mice, indicating that CHIKV can cause bone pathologies (11, 12). In another study, mice infected with a similar arthritogenic alphavirus, Ross River virus (RRV) resulted in significant bone loss. (13). In humans, magnetic resonance imaging (MRI) studies revealed that CHIKV infection is associated with erosive arthritis (14, 15). Taken together, these studies suggest alphavirus infection can affect bone homeostasis and thus contribute to arthritic-like conditions.

Mesenchymal stem cells (MSCs) are multipotent, non-hematopoietic stromal cells which can self-renew and differentiate into various cell lineages (16). MSCs can be derived from umbilical cord blood, adipose tissue and bone marrow (16). Bone marrow-derived MSCs (BMMSCs) have trilineage differentiation potential and they can differentiate into osteogenic, chondrogenic or adipogenic cell lineage (17). The osteogenic differentiation of BMMSCs is important for bone-remodeling, and the inability of BMMSCs to differentiate into the osteogenic lineage may lead to an imbalance in bone remodeling and eventually bone loss (18–20). A few studies have shown that virus infection of BMMSCs can affect the properties and function of these cells (21, 22).

In this study, we investigated the susceptibility and response of BMMSCs to CHIKV infection. We hypothesized that CHIKV can infect BMMSCs and affect the osteogenic differentiation of BMMSCs. Our results show that CHIKV can productively infect BMMSCs. Importantly, we observed that viral infection significantly impaired the function of the osteogenic cells, as evidenced by the decrease in alkaline phosphatase (ALP) activity at 14 days post infection (dpi) and rate of mineralization at 7 and 14 dpi compared to mock-infected control. Together, these findings indicate CHIKV can infect BMMSCs and disrupt BMMSC-derived osteogenic cell function.

## Results

### BMMSCs are permissive to CHIKV infection

CHIKV infection has been associated with bone pathology, implying its role in disruption of bone homeostasis (11, 12, 14, 15). BMMSC-derived osteogenic differentiation is essential for bone homeostasis (18–20). Recent studies show that viral infection can affect the function of BMMSCs and BMMSC-derived osteogenic cells (21, 22). However, to date, it is unknown whether alphaviruses can infect BMMSCs and disrupt osteogenic cell function. Permissivity of BMMSCs to CHIKV infection was determined by infecting cells under acute infection condition (Fig.1). To detect the presence of infection in BMMSCs, immunofluorescence assays (IFA) were performed at 24 hours post-infection (hpi). Infection was confirmed by visualizing the presence of viral non-structural protein 4 (nsP4) in infected cells at all MOIs tested (Fig. 2A). To detect replication of virus in infected cells, the expression of viral non-structural protein 1 (nsP1) gene was quantified by qRT-PCR. Infected cells showed a significant increase in nsP1 gene expression at 24 hpi with increased MOIs (Fig. 2B). Morphological analysis of infected cultures at 48 hpi showed the presence of cytopathic effect (CPE), particularly at higher MOIs (Fig. S1A). CPE was quantified at 48 hpi using Viral ToxGlo assay by determining ATP content. Higher MOIs (0.1 and 1.0) resulted in significant CPE as evidenced by the decrease in luminescence signal (Fig. S1B). Lower MOIs (0.001 and 0.01) resulted in productive viral infection as evidenced by IFA and qRT-PCR with minimal CPE. Collectively, these data indicate that BMMSCs are susceptible to CHIKV infection.

**Fig 1.**
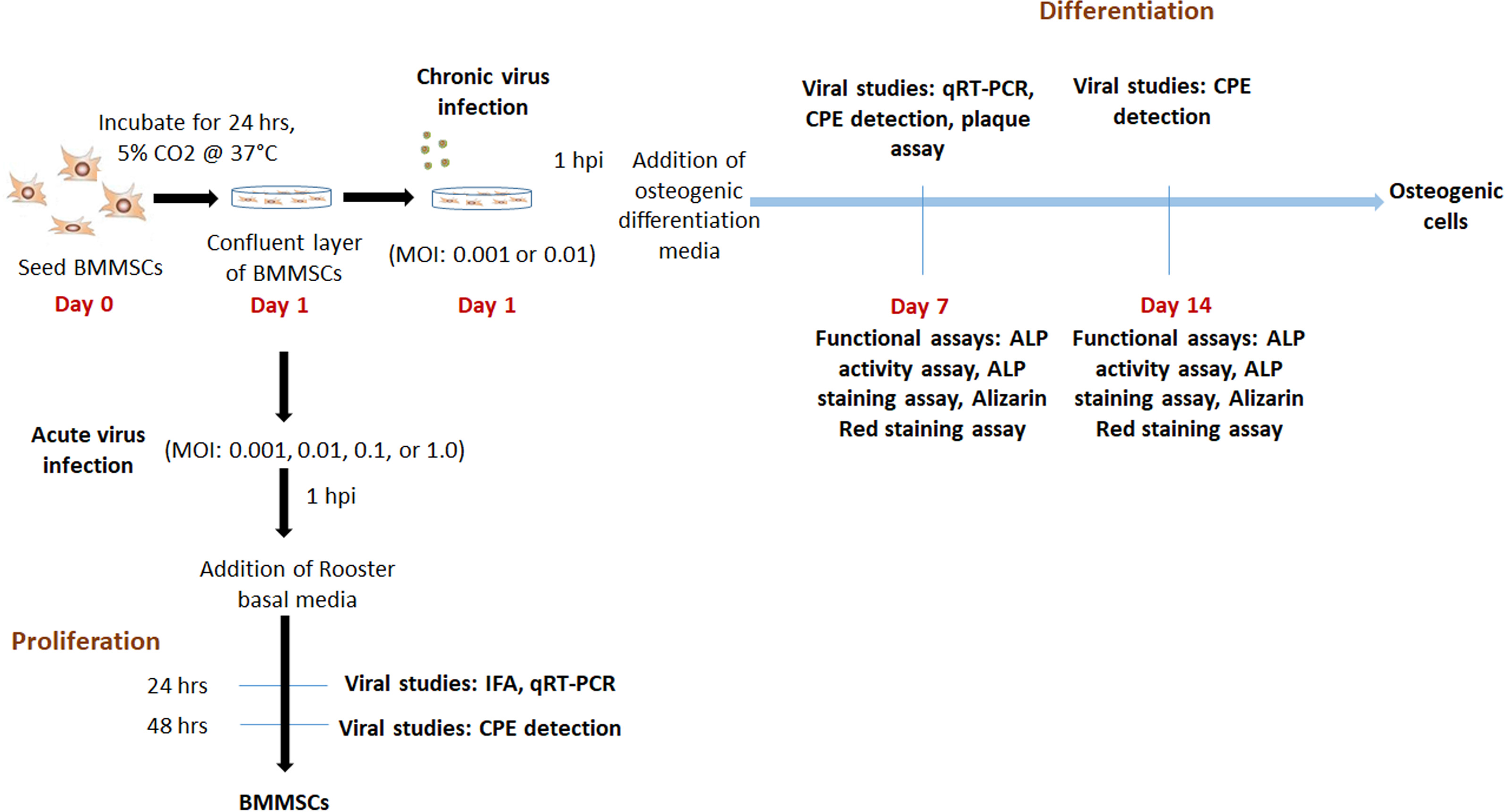
Experimental outline.

**Fig 2.**
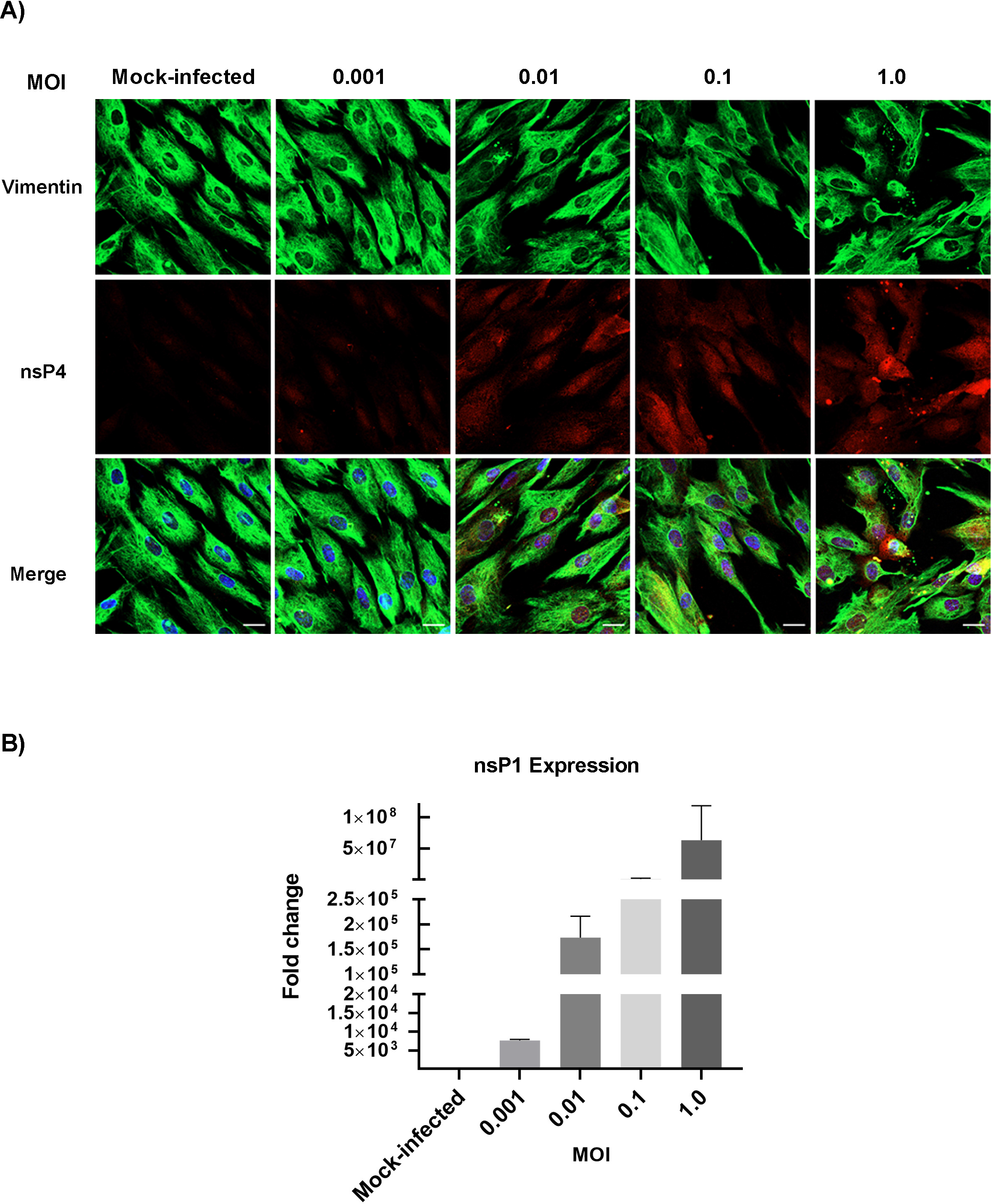
CHIKV infects and replicates in BMMSCs. BMMSCs were mock-infected or infected with CHIKV under acute viral infection: MOIs of 0.001, 0.01, 0.1 or 1.0 (A) Representative immunofluorescence images of cells fixed and immunostained at 24 hpi with antibodies against the viral nsP4 protein (red), Vimentin (green), and Hoechst nuclear stain (blue). Scale bar=20μm. (B) Viral nsP1 gene expression was quantified by qRT-PCR at 24 hpi. Bar graph shows fold change in gene expression. Fold change was calculated by 2^-ΔΔCt method. Error bars show ±SEM.

### CHIKV infection interferes with function of osteogenic cells

#### BMMSC-derived osteogenic cells are susceptible to CHIKV infection

Osteogenic differentiation of BMMSCs plays an important role in bone homeostasis (18–20). In addition, viral infection have been associated with altered function of BMMSC-derived osteogenic cells (21). Therefore, we next investigated the fate of CHIKV-infected, differentiated BMMSCs. Based on our initial data we selected a MOI of 0.001 and 0.01 for further “chronic infection” experiments. Cells were infected and differentiated as described in Figure 1. Osteogenic differentiation was confirmed by Alizarin Red staining assay. This assay detects the deposition of calcium phosphate crystals, formed during matrix mineralization by osteogenic cells (23). Presence of red stained calcium nodes was clearly detected in differentiated cells at 7 and 14 days post differentiation (dpd) compared to undifferentiated controls (Fig 3). To determine the susceptibility of the osteogenic cells to CHIKV infection, viral replication was confirmed by nsP1 gene expression using qRT-PCR at 7 dpi. A significant increase in nsP1 expression was observed in infected cells compared to mock (Fig. 3B). Next, we confirmed the production of infectious virus particles at 7 dpi by plaque assay (Fig. 3C). Similarly, we observed an increase in infectious virus particle production. We next determined the viability of the osteogenic cells during infection. Brightfield images of infected cells at 7 dpi showed evidence of CPE, particularly at MOI 0.01 (Fig. 3D). Interestingly, CPE quantification done by Viral ToxGlo assay resulted in minimal CPE production when compared to mock-infected cells at 7 dpi (Fig. S2A). Morphological analysis and CPE quantification at 14 dpi showed similar results (Fig. S2B and C). Taken together, these results demonstrate that osteogenic cells are susceptible to CHIKV infection and results in minimal CPE at low MOIs.

**Fig 3.**
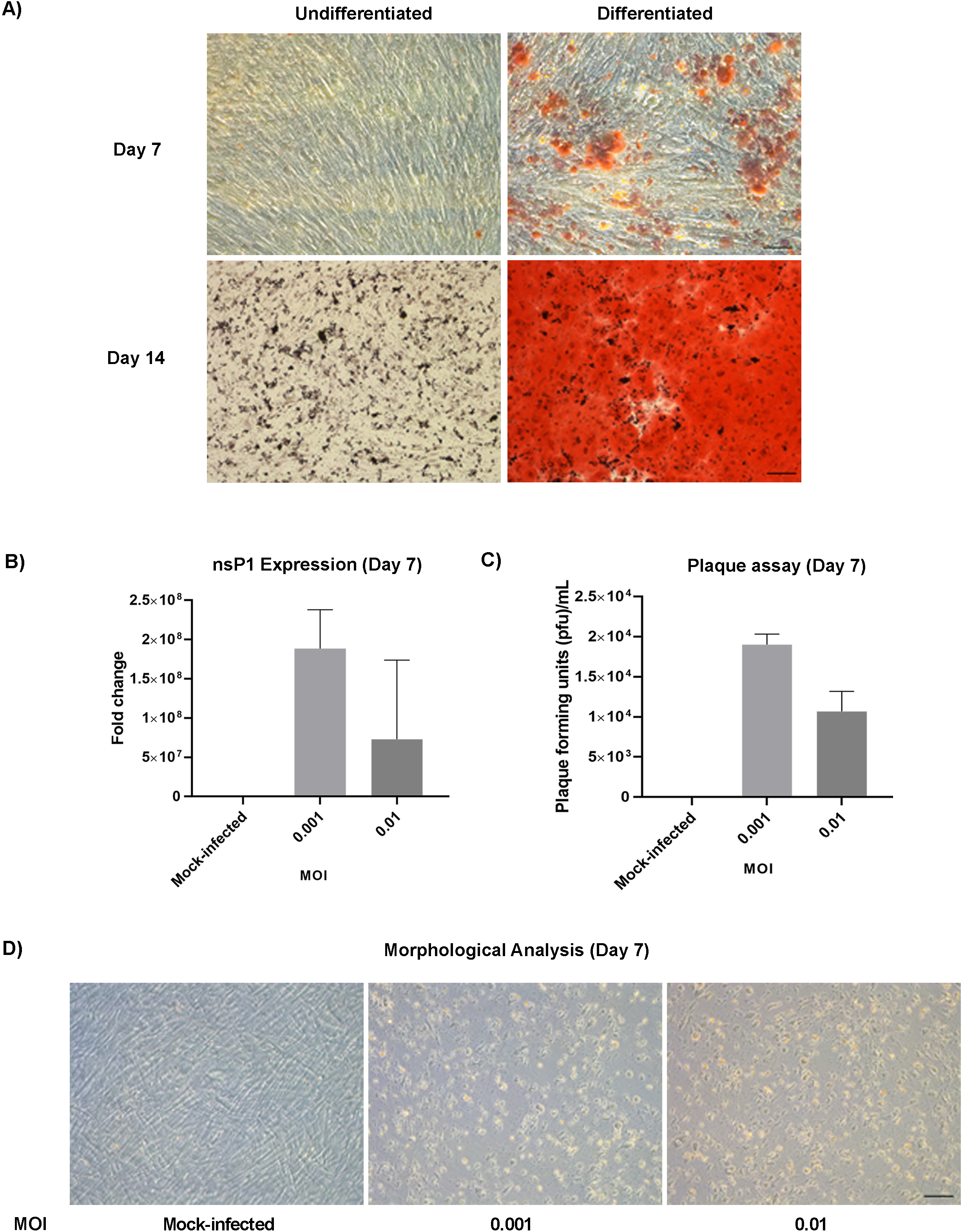
Osteogenic differentiation of BMMSCs and susceptibility of osteogenic cells to CHIKV infection. (A) Representative brightfield images of mock infected, differentiated and undifferentiated BMMSCs, stained with Alizarin Red at 7 and 14 dpd. Scale bar=0.2μm. (B) Viral nsP1 gene expression was quantified by qRT-PCR at 7 dpi. Bar graph shows fold change in gene expression in infected cells compared to mock-infected control. Fold change was calculated by 2^-ΔΔCt method. n=3. (C) Plaque assays were performed using culture supernatant of infected BMMSCs. n=3. Error bars show ±SEM. (D) Representative bright field images of morphological analysis of BMMSCs at 7 dpi to detect CPE. n=3. Scale bar=0.2 μm.

#### CHIKV infection impairs function of osteogenic cells

Prior studies have shown viral infection of osteogenic cells causes impaired ALP activity during differentiation (21). To examine the effect of CHIKV infection on ALP activity, BMMSCs were infected as previously outlined (Fig 1). ALP activity and ALP staining assays were performed. No significant change in ALP activity was observed in infected cells at 7 dpi (Fig. 4A). However, a significant reduction in the ALP activity was observed in infected cells at 14 dpi (Fig. 4B). The infected cells resulted in reduced ALP staining intensity at both 7 and 14 dpi (Fig. 4C). Next we performed Alizarin Red staining assay to detect the effect of viral infection on deposition of calcium phosphate crystals. BMMSC were infected as described in Figure. 1 and Alizarin Red staining assay was done at 7 and 14 dpi. A significant decrease in calcium phosphate deposits was observed in infected cells at 7 and 14 dpi (Fig. 4D). Together, these results demonstrate for the first time that CHIKV infection impairs the function of BMMSC-derived osteogenic cells.

**Fig 4.**
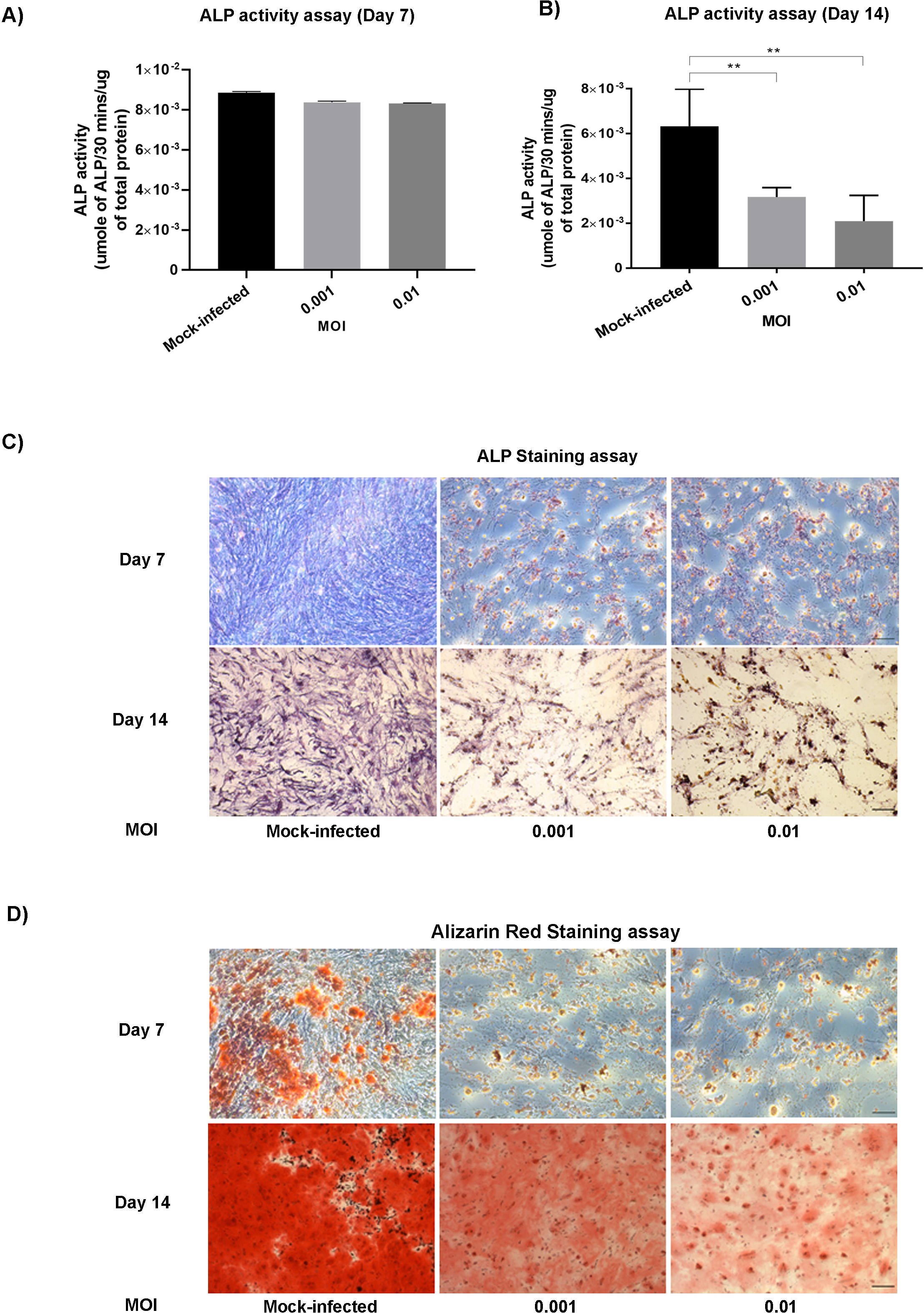
Effect of CHIKV infection on osteogenic cells. BMMSCs were mock-infected or infected with CHIKV under chronic viral infection at MOIs of 0.001 or 0.01. (A and B) Bar graphs show intracellular ALP activity detected by ALP activity assay at 7 and 14 dpi. ALP levels were normalized against total protein. n=5. Error bars show ±SEM. Significant changes are represented as p-values (**p<0.001) (C) Representative bright field images of infected and mock infected BMMSCs stained by ALP staining assay to detect extracellular ALP activity at 7 and 14 dpi. (D) Representative brightfield images of infected and mock infected BMMSCs, stained with Alizarin Red at 7 and 14 dpi. Scale bar=0.2 μm.

## Discussion

Joint pathology and arthritic-like conditions are associated with CHIKV infection, however the mechanism underlying these conditions remains poorly defined (11, 12, 14, 15, 24). BMMSCs are non-hematopoietic multipotent stem cells which can differentiate into adipogenic, osteogenic, and chondrogenic lineages (16, 17). These cells can secrete and respond to different signaling molecules and undergo osteogenic differentiation thereby playing an essential role in bone homeostasis (18, 25). Previous studies have shown that virus infection can affect functions of BMMSCs (21, 22, 26), but it is unclear whether BMMSCs are susceptible to alphavirus infection.

Osteogenic cells will terminally differentiate into osteoblasts (18, 23). A prior report demonstrated that osteoblasts are susceptible to CHIKV infection (27). However, it is unclear whether osteoblast progenitor cells are susceptible to infection. Moreover, whether infection affects differentiation and/or function. The results of our study demonstrate that both BMMSCs and BMMSC-derived osteogenic cells are susceptible to CHIKV infection. Prior studies have shown that viral infection of BMMSCs can alter function, therefore we hypothesize that ability of CHIKV to infect these cells will similarly affect function (22). In our preliminary experiments, we observed CPE during infection of both BMMSCs and osteogenic cells at relatively high MOI. However, minimal CPE was observed at low MOI. We therefore selected lower MOI infections for our chronic infection experiments.

Previous studies demonstrated that viral infection causes impaired matrix mineralization and ALP activity (21, 22). Our study revealed that despite a lack of significant change in ALP activity at 7 dpi, there was a significant reduction in ALP activity at 14 dpi. Moreover, reduction in ALP staining intensity was observed at both 7 and 14 dpi. The results obtained are similar to a prior study showing the effect of viral infection on ALP activity (21). The lack of significant change in ALP activity at 7 dpi was likely due the observed absence of a difference in ALP activity between undifferentiated and differentiated osteogenic cells at 7 dpd (data shown). Calcium phosphate deposition due to mineralization is a hallmark of osteogenic differentiation (23, 28). We found that at 7 and 14 dpi, CHIKV caused a significant reduction in calcium phosphate deposition in infected cells. This indicates that CHIKV infection disrupts the mineralization function of osteogenic cells, in that way likely disrupts bone homeostasis. It is important to note that matrix mineralization is also associated with the production of extracellular matrix (ECM) proteins including fibronectin, vitronectin, laminin, osteopontin, and osteonectin (23, 28–30). Thus, future studies will be aimed at examining the expression and production of these proteins in the context of viral infection.

*In vivo* differentiation of BMMSCs into the osteogenic cell lineage depends on their interaction with other cells in the joint space (23, 31–33). Hence, co-culturing of BMMSCs with cells, including osteoclasts, synovial fibroblasts, macrophages and chondrocytes in the presence of CHIKV will lead to a better understanding the mechanism underlying the disruption of bone homeostasis (34–36).

It is well established that the receptor activator of nuclear factor kappa-Β ligand (RANKL)/osteoprotegerin (OPG) ratio plays an important role in bone homeostasis and drives osteoclastogenesis (37–40), the dysregulation of which likely contributes to CHIKV-induced bone pathology. It has been reported that CHIKV infection of osteoblasts can alter RANKL/OPG ratio (41). Based on our findings, it will be interesting to examine the contribution of CHIKV infection of osteoblast progenitors *in vitro* and *in vivo* and the effect on RANKL/OPG levels.

In our study, we report that BMMSCs are susceptible to CHIKV infection, and infection impairs ALP activity and calcium phosphate deposition in osteogenic cells. Our results indicate that CHIKV infection leads to functionally altered osteogenic cells and likely leads to dysregulation in bone homeostasis. Thus, studying BMMSCs in the context of CHIKV infection can serve as an appropriate model for better understanding viral-induced bone pathology.

## Materials and Methods

### Cell culture, Compounds and Virus

Human BMMSCs were purchased from Roosterbio (USA). CHIKV (181/25 clone) was purchased from BEI Resources (USA).

BMMSCs were infected at MOI 0.001, 0.01, 0.1 or 1 and incubated for 1 h at 37°C, 5% CO_2_. At one-hour post infection (hpi), cells were washed with DPBS (Corning, USA), replenished with Rooster Basal MSC (RBM) media (RoosterBio, USA) and incubated at 37°C, 5% CO_2_ for 2 days for proliferation. For the purposes of this study, the 2-day incubation period was considered an acute infection (Fig. 1).

For differentiation study, BMMSCs were infected at MOI 0.001 or 0.01 and incubated for 1 h at 37°C, 5% CO_2_. After 1 hpi, cells were washed with DPBS, replenished with osteogenic differentiation media containing mesenchymal stem cell expansion medium (Millipore, USA) and supplemented with 10 mM beta glycerophosphate (Sigma, USA), 0.1 μM dexamethasone (Sigma, USA) and 200 μM ascorbic acid (Sigma, USA). Media change was done every 2 dpi. For this study, this was considered as a chronic infection (Fig. 1). For these experiments, mock-infected undifferentiated cells were negative controls, while mock-infected differentiated cells were positive controls.

Baby hamster kidney cells (BHK-21; ATCC CCL10) were cultured in Dulbecco’s modified Eagle’s medium (DMEM) (Invitrogen, USA) and supplemented with 10% Fetal bovine serum (FBS) (Gibco, USA)

### Determination of CPE

Cells at a density of 1×10^5^ per well were seeded in 6-well plates and incubated for 24 h, at 37°C. A confluent monolayer of cells were infected under both acute and chronic conditions as mentioned above. To determine CPE, infected cells were observed using the DM1/MC120 bright field microscope (Leica, Germany). Images were collected every 24 h for acute infection and at 7 and 14 dpi for chronic infection.

The viral CPE was quantified by the Viral ToxGlo assay (Promega, USA). This assay detects the amount of ATP produced by the cells (42). CPE produced due to viral infection leads to decreased ATP production which can be quantified. Cells at a density of 3×10^3^ per well were seeded in a 96-well plate and incubated for 24 h at 37°C. For quantification of CPE in acute infection, Viral Tox Glo assay was performed at 48 hpi, and for chronic infection, the assay was performed at 7 and 14 dpi, according to manufacturer’s instructions. The change in ATP production in infected versus mock-infected BMMSCs was measured using a Tecan Spark® microplate luminometer (Tecan Trading AG, Switzerland).

### Immunofluorescence assay

Cells at a density of 2.5×10^4^ per well were seeded in 4-well plate containing glass cover slips in each well and incubated for 24 h, at 37 °C. Mock or virus-infected cells were fixed in 4% paraformaldehyde (PFA) (Electron Microscopy Sciences, USA) for 30 min and permeabilized with 0.1% Triton X (Fisher Bioreagents, USA) for 10 mins. Blocking was done in 3% Bovine Serum Albumin-Phosphate Buffered Saline (PBS/BSA) for 1 h. Virus was stained with antibody against CHIKV non-structural protein (nsP4) (kindly provided by Dr. Andres Merits). BMMSCs were stained with mouse anti-Vimentin antibody (V9) (Life Technologies, USA). Following overnight incubation with primary antibody at 4°C, the cells were stained with Alexa-Fluor 568 goat anti-rabbit IgG and Alexa Fluor 488 donkey anti-mouse IgG (Life Technologies, USA) for 1 h at room temperature. Nuclear staining was done with Hoechst 33642 (Invitrogen, USA) for 30 min. Images were taken and processed using Zeiss LSM 800 with Airyscan (Germany).

### ALP activity and ALP staining assay

ALP enzyme activity was determined by ALP activity assay (43). Cells at a density of 2.5×10^4^ per well were seeded in a 24-well plate and incubated for 24 h at 37°C. Cells were then infected as aforementioned. At 7 and 14 dpi, the infected cells were lysed by freeze-thaw method in a buffer containing Triton X-100 (0.1% v/v) (Fischer Scientific, USA), 1 mM MgCl_2_ (Alfa Aesar, USA), 20 mM Tris (Fischer Scientific, USA). The cell lysate was used to perform ALP activity assays using the ALP activity kit (Sigma-Aldrich, USA) according to the manufacturer’s instructions. The ALP activity assay uses p-nitrophenyl phosphate (pNPP) as substrate which is dephosphorylated by the ALP enzyme forming *p*-nitrophenol which was measured spectrophotometrically at 405 nm in Synergy H1 Hybrid Reader (BioTek, USA) (43). The total protein content in the infected cells was determined using a BCA protein assay kit (Thermo Scientific, USA) with bovine serum albumin as a standard. The ALP activity was expressed as micromole of p-nitrophenol formed per 30 minutes per microgram of total protein (μmol per 30 min per μg protein).

Extracellular ALP expression was examined by ALP staining assay (44). Cells at a density of 3×10^4^ per well were seeded in a 24-well plate and incubated for 24 h at 37°C. At 7 and 14 dpi, ALP staining was performed using ALP leukocyte kit (Sigma-Aldrich, USA) according to manufacturer’s instructions. Images were taken using a DM1/MC120 microscope (Leica, Germany) at 20× magnification.

### Alizarin Red staining

Cells at a density of 5×10^4^ per well were seeded in a 24-well plate and incubated for 24 h at 37°C. At 7 and 14 dpi, Alizarin Red staining assay was performed using Alizarin Red solution (EMD Millipore, Germany) according to manufacturer’s instructions. Alizarin Red stains the calcium phosphate deposits formed by matrix mineralization during osteogenic differentiation (23). All images were taken using DM1/MC120 microscope (Leica, Germany) at 20× magnification.

### Quantitative RT-PCR (qRT-PCR)

Cells at a density of 1×10^5^ per well were seeded in a 6-well plate and incubated for 24 h at 37°C. For acute and chronic infection, cells were collected at 24 hpi and 7 dpi respectively, and lysed in RLT buffer (Qiagen, Germany) for RNA isolation. RNA isolation was performed using RNeasy Mini kit (Qiagen, Germany) according to the manufacturer’s instructions. RNA was quantified and total RNA was reverse transcribed using a qScript cDNA synthesis kit (Quantabio, USA) according to the manufacturer’s instructions. SYBR Green Real-Time PCR was performed on a StepOnePlus Real-Time PCR System (Thermo Scientific, USA) using SSO Advanced SYBR Green Supermix (Bio-Rad, USA) and primer for the nsP1 gene (Applied Biosystems, USA) (5’-GGGCTATTCTCTAAACCGTTGGT-3’ and 5’-CTCCCGGCCTATTATCCCAAT-3’) according to the manufacturer’s instructions with the following conditions: (i) PCR initial activation step, 95°C for 3 min, 1 cycle, and (ii) three-step cycling, 95°C for 15s, followed by 60°C for 1min and 95°C for 15s, 40 cycles. The fold change in mRNA expression relative to mock-infected samples was calculated using the 2^-ΔΔCt method. Transcript levels were normalized using Hypoxanthine phosphoribosyltransferase 1 (HPRT1) (Hs02800695_m1) (Thermo Scientific, USA)

### Plaque assay

BMMSCs at a density of 1×10^5^ per well were seeded in a 6-well plate and incubated for 24 h, at 37°C. The cell culture supernatants were collected from infected and mock-infected cells at 7dpi and after serial dilution, were used to perform plaque assay. BHK21 cells were seeded into 24-well plates at a density of 1×10^5^ cells per well and cultured to confluence. Confluent layer of BHK21 cells were infected in triplicate with each dilution of the culture supernatant of infected BMMSCs and incubated for 1 h and 15 min at 37°C, 5% CO_2_. Cells were then overlaid with DMEM containing 5% FBS and 0.6% agarose (Fischer scientific, USA) and incubated at 37°C for 48 h. After 48 h, the agarose plugs were removed, and the cells were fixed and stained with 0.1% crystal violet solution containing 3.2% PFA. The plaques were counted, and the viral titers were expressed as plaque forming units/mL (PFU/mL).

## Statistical analysis

All statistical analyses were performed using GraphPad Prism 8. Data were represented as the mean ± standard error of the mean (SEM). Significant differences between the experimental groups were determined using Student’s t test. P values < 0.001 were considered significant.

## Acknowledgments

We thank Dr. Leah Cook for critical reading of the manuscript and valuable suggestions. We thank Janice A. Taylor and James R. Talaska of the Advanced Microscopy Core Facility at the University of Nebraska Medical Center for providing assistance with confocal microscopy. We also thank Dr. Andres Merits for kindly providing the nsP4 antibody. This study was supported by NIH/NIAID 1R21AI140026-01 (SPR & BD) and start-up funds (SPR).

## Figure Legends

Fig. S1. CHIKV infection produces CPE in BMMSCs. BMMSCs were infected at MOIs of 0.001, 0.01, 0.1 or 1.0. (A) Representative brightfield images of morphological analysis of BMMSCs at 48 hpi. Scale bar=0.2 μm. (B) Quantification of CPE was performed at 48 hpi using Viral ToxGlo assay. n=3. Error bars show ±SEM.

Fig. S2. CHIKV produces CPE in osteogenic cells. BMMSCs were mock-infected or infected with CHIKV under chronic viral infection: MOIs of 0.001 or 0.01. (A) Quantification of CPE was performed at 7 dpi using Viral ToxGlo assay. n=3. Error bars show ±SEM.

(B) Representative brightfield images of morphological analysis of BMMSCs at 14 dpi. Scale bar=0.2 μm. (C) Quantification of CPE was performed at 14 dpi using Viral ToxGlo assay. n=3. Error bars show ±SEM.

